# The general anesthetic isoflurane inhibits calcium activity in cerebrovascular endothelial cells and disrupts vascular tone

**DOI:** 10.1101/2022.03.25.485881

**Authors:** Lingyan Shi, Adrián Rodríguez-Contreras

## Abstract

Calcium signaling in cerebrovascular endothelial cells (CVECs) has been identified to play key physiological and pathological roles in blood brain barrier function and neurovascular coupling, which involve dynamic changes in vessel diameter. However, there are no studies that measured correlated changes in vessel diameter and calcium activity in CVECs in vivo. In this study, we used the general anesthetic isoflurane (ISO) to induce a maximally dilated state in cortical blood vessels and measured the effects of the manipulation on CVEC calcium reporter activity in awake Cdh5^BAC^-GCaMP mice by use of two-photon fluorescence microscopy through thinned skull cranial windows. For the first time, we report dual effects of ISO on calcium activity in cerebral blood vessels of different diameter. During anesthesia induction ISO exposure triggered a short latency synchronous increase in calcium activity, followed by a period of activity suppression in small, medium, and large diameter vessels. Furthermore, during anesthesia maintenance calcium activity was desynchronized, and the relationship between vascular tone and calcium activity was disrupted in all vessel types. Based on these results we propose that there is a feedback mechanism between intracellular calcium fluctuations in CVECs and the maintenance of cerebrovascular tone.

## Introduction

Vascular endothelial cells (VECs) are involved in the regulation of vascular tone, inflammation, coagulation, growth and permeability (Andresen et al., 2006; Filippini et al. 2019). The vascular endothelium produces a number of vasodilators such as nitric oxide (NO), prostanoids, endothelial-derived hyperpolarizing factor, and vasoconstrictors such as endothelin. In the vascular endothelium, NO is produced by endothelial nitric oxide synthase (eNOS), which is regulated by calcium. Constitutive release of NO regulates the basal vascular tone that maintains the vasculature in a vasodilated state. Many factors control eNOS and NO production by regulating intracellular calcium level (Andresen et al., 2006). Although major progress in understanding the spatial and temporal properties of calcium signaling in VECs has been achieved using isolated arteries and synthetic calcium indicators (Mumtaz et al., 2011; Taylor and Francis, 2014; Kansui et al., 2008; Bagher et al., 2012; Dora and Hill, 2013), it has been more difficult to study VEC calcium signaling in *in vivo* conditions.

Development of genetically encoded calcium indicators expressed in VECs has been used to show that agonist-induced increases in intracellular calcium cause vasodilation of peripheral arteries *in vivo* (Tallini et al., 2007), and more recently, it was used to measure correlated changes in calcium activity during trans-endothelial leukocyte migration (Dalal et al., 2021). However, these studies used anesthetized animals and the extent to which anesthesia affected the relationship between intracellular calcium in VECs and other processes of interest remains unknown. This issue is important since the functional properties of VECs may vary notably amongst different organs, including the brain, where an increase in neuronal activity leads to a local elevation in cerebral blood flow mediated by vasoactive compounds that act on cerebral blood vessels (Filippini et al., 2019; Guerra et al., 2018; Thakore and Earley, 2019). Furthermore, general anesthetics such as isoflurane (ISO) have been shown to cause large disruptions in cerebral blood flow (Li et al., 2014; Shumkova et al., 2021), brain metabolism (Slupe and Kirsch, 2018), neurovascular coupling (Gao et al., 2016; Schlegel et al., 2015), and functional connectivity (Ciobanu et al., 2012; Constantinides and Murphy, 2016; Bukhari et al., 2018; Stenroos et al., 2021). One of the most recent studies used multi-exposure speckle imaging to examine the effects of ISO on the dynamics of the cerebral vasculature and blood flow in awake mice. Dramatic vasodilation and blood flow increase were measured within minutes after ISO induction (Sullender et al., 2022). These observations raise the possibility that general anesthetics may have direct and indirect effects on the physiology of cerebrovascular endothelial cells (CVECs). However, to our knowledge, no study has directly monitored CVEC calcium activity, vascular tone and their responses to general anesthesia from an awake state in vivo.

In this study, we used ISO to induce a maximally dilated state in cortical blood vessels and calcium reporter activity in CVECs was imaged with two-photon fluorescence microscopy through thinned skull windows before, during, and after ISO exposure in awake Cdh5^BAC^-GCaMP mice (Figure 1). We report for the first time that ISO has dual effects on calcium reporter activity in cerebral blood vessels of different diameter. During anesthesia induction, ISO exposure triggered an increase in CVEC calcium reporter activity that was correlated with vasodilation in medium and large diameter vessels but not in small diameter vessels. Sustained ISO exposure during anesthesia maintenance caused an inhibition of calcium activity in all vessel types examined. ISO exposure also disrupted the relationship between vessel tone and calcium activity in small, medium and large vessels, despite their different vasodilation properties. After ISO was removed, the calcium-vessel tone relationship remained disrupted. We propose that there is a feedback mechanism between intracellular calcium fluctuations in CVECs and the maintenance of cerebrovascular tone.

**Figure 1.**
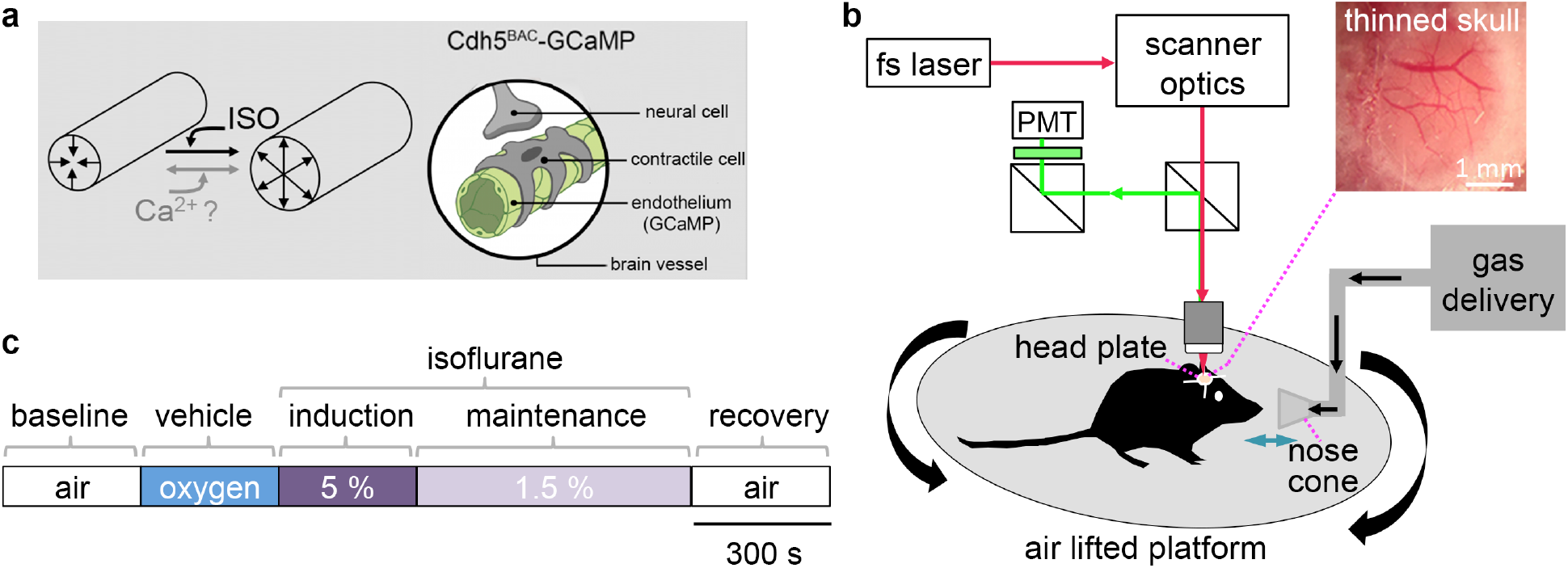
Research question and experimental approach. **a**, The relationship between changes in the second messenger calcium and vascular tone was investigated by manipulating vasodilation with the general anesthetic isoflurane (ISO). ISO was used to dilate cerebral blood vessels and determine the effects on calcium signaling in vascular endothelial cells of Cdh5^BAC^-GCaMP mice. **b**, Two-photon fluorescence microscope setup used in this study. A femtosecond (fs) laser was used to excite the genetically encoded calcium sensor GCaMP and emitted fluorescence was point scanned to a photomultiplier tube (PMT) for time-lapse image acquisition. Inset shows an image of the thinned skull window. A retractable nose cone was used to deliver ISO to a head restrained mouse from either sex. **c**, Protocol for gas exposure included baseline, vehicle, induction, maintenance and recovery conditions.

## Materials and Methods

### Animals

All experiments and procedures were approved by the City College of New York Institutional Animal Care and Use Committee (IACUC). Breeding pairs from the Cdh5^BAC^-GCaMP mouse strain were obtained from CHROMus™ (Cornell University), and are also available from Jackson Laboratories (JAX033342). Mice from the Cdh5^BAC^-GCaMP strain have constitutive expression of the genetically encoded calcium indicator GCaMP8 by insertion of a GCaMP8 cassette in the calcium-dependent cell adhesion molecule cadherin 5 locus (Cdh5 or VE-cadherin), which is active in endothelial cells and myeloid cells (Vestweber, 2008; Harris and Nelson, 2010). Breeding pairs of wild type and heterozygous Cdh5^BAC^-GCaMP mice were maintained with free access to food and water in climate-controlled rooms with a 12 hr light/dark cycle (lights on at 6 am). Mouse litters were housed with their parents until weaned at postnatal day 19 (P19). After weaning, mice of either sex were genotyped and housed in same sex pairs or trios until used for experiments. Experiments were not designed to evaluate differences between sexes, so data from females and males was pooled for statistical analysis.

### Thinned skull surgery and recovery

Cranial imaging windows were prepared following standard techniques. A total of ten mice between ages P49 to P83 were processed for surgery. Animals were anesthetized with isoflurane (ISO) mixed in oxygen vehicle at 5% for induction and 1.5% for maintenance. Mice were then head fixed in a mouse stereotaxic. After removal of the scalp, a circular skull region over the somato-sensory cortex (AP: -4.5, ML: +2.0) was thinned using a ceramic piece attached to a high-speed dental drill. A circular glass coverslip (3 mm diameter, Warner) was secured over the polished skull area using cyanoacrylate glue (Loctite 401). A four-point circular head plate (NeuroTar) to be used for head restraint was secured to the skull around the coverslip using cyanoacrylate glue and dental cement. Animals were allowed to recover individually on cages warmed by a heating blanket for up to an hour before being placed back with cagemates. Post-surgery recovery lasted between 6 and 46 days. One female and two males showed visible signs of trauma one-week post surgery and were removed from the study. Approximately one-week before imaging experiments, animals were habituated to handling (days 1 to 3), and to head restraint on the air lifted platform used for two-photon fluorescence imaging (Neurotar; days 4-7).

### Two-photon fluorescence imaging

Two-photon fluorescence imaging was performed with an Ultima 2P In Vivo microscope equipped with galvanometer scanning mirrors (Bruker). A Chameleon laser (Coherent) was tuned to 920 nm and kept below 50 mW output measured before the objective. GCaMP8 fluorescence was passed through a 520/15 filter (Chroma) and placed in front of a photomultiplier (PMT) detector (Figure 1b). Images were digitized at 16-bit depth and acquired at 1 Hz frame rate and 512×512 pixel resolution using 20x/0.6 or 40x/0.8 water immersion objectives (Olympus). Prior to imaging, mice were allowed to acclimate in the head restraint of the air lifted platform for at least 10 min. Time-lapse imaging was done for 25-30 min at depths between 100 to 250 μm below the thinned skull surface following the sequence pictured in Figure 1c. First, awake air breathing mice were imaged for 5 min to obtain baseline data. A nose cone attached to a flexible arm was then placed in front of the mouse snout, and the mouse was imaged for 5 minutes during exposure to oxygen vehicle delivered at 1L min^-1^. Next, mice were imaged for 5 min during induction with 5% ISO mixed with oxygen, followed by 5 or 10 minutes of maintenance in 1.5% ISO mixed with oxygen. At the end of gas exposure, the nose cone was removed and the mouse was allowed to breathe air for 5 minutes while imaging calcium reporter activity. During anesthetic exposure, body temperature was maintained with a heating pad set at 37°C (Physitemp), or with a disposable heating pad at 25-30°C (Hot Hands). After imaging experiments animals were placed in a warm cage until fully recovered and returned to their home cage.

### Image processing and analysis

All images were analyzed offline with Fiji (Image J version 2.3.0/1.53f, National Institutes of Health). Time-lapse sequences were denoised using the despeckle function and converted from 16-bit to 8-bit image depth. Line and region of interest (ROI) measurements were collected semi-automatically with the multi measure function in the ROI manager. The line tool was used to measure vessel diameter in vessel regions that showed changes in fluorescence over time. Average vessel diameter was obtained from 60 consecutive replicates per vessel at the end of the baseline, induction and recovery conditions. Time course of vessel diameter was obtained from 120 consecutive measurements at the start of the induction condition. ROI were drawn with the box or the polygon line tools around vessel segments of interest. Distance, area and integrated fluorescence intensity values were saved in csv files and processed in Igor Pro 8 (Wavemetrics) or Prism 6 (GraphPad).

Integrated fluorescence recordings from box or polygon ROI were examined in Igor Pro software using the Neuromatic toolbox (Rothman and Silver, 2018). DF/F_0_ values were calculated according to equation:

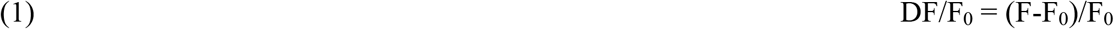

where F_0_ represents the mean baseline fluorescence, obtained as the average fluorescence from 20 s to 50 s windows of quiescent activity. A calcium reporter fluorescence event F was considered to occur at a threshold of three standard deviations above F_0_.

Binned normalized DF/F_0_ (_N_DF/F_0_) recordings were fitted to the sigmoid equation:

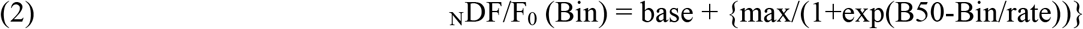

where base is the minimum _N_DF/F_0_ = 0; max is the maximum _N_DF/F_0_ = 1; B50 is the bin at which the _N_DF/F_0_ reached 50% of its maximum; and rate represents the change of _N_DF/F_0_ over time (Bin width = 50 s).

Integrated fluorescence recordings from polygon ROI were correlated with the corresponding polygon ROI area. Measurements were obtained every 10^th^ frame for a total of 30 replicates per vessel per condition and processed to generate DF/F_mean_ and DA/A_mean_ traces according to equation:

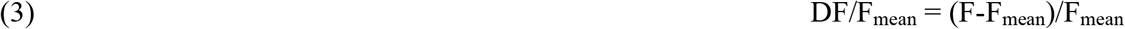

where F_mean_ represents the mean fluorescence obtained as the average fluorescence from all 30 replicates, and equation:

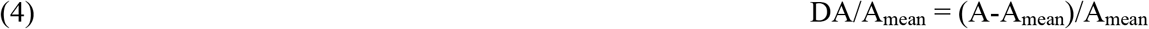

where A_mean_ represents the mean area obtained as the average area from all 30 replicates.

Although DA/A_mean_ vs DF/F_mean_ scatter plots were generated to visualize correlated changes between calcium reporter fluorescence and vessel area, correlation coefficients were formally obtained using Prism statistical software.

### Statistics

Statistical analyses were performed with Prism 6. The D’Agostino and Pearson omnibus normality test was used to choose between a parametric or a non-parametric test, and statistical significance was set at alpha = 0.05. For multiple comparisons, the Kruskal-Wallis test was used and *p* values were corrected with the Dunn’s test. A two-tailed nonparametric Spearman correlation was performed for each DA/A_mean_ and DF/F_mean_ data pair with a 95% confidence interval and 4 significant digits. Unless otherwise stated, data represent mean ± SD.

## Results

### Isoflurane (ISO) dilates vessels of medium and large diameter, but not vessels of small diameter

We imaged fourteen cortical blood vessels of different diameter. First, we determined the effect of ISO on mean vessel diameter by comparing baseline, anesthesia induction and recovery conditions. Blood vessels were grouped into small, medium or large diameter groups. The mean diameter of small vessels was 8.3 ± 1.5 μm in baseline, 8.1 ± 1.3 μm in induction, and 8.1 ± 4.4 μm in recovery conditions. These values were not statistically different from each other (**Figure 2a**; n = 4 vessels from 2 mice; Kruskal-Wallis test (3, 2.901) and Dunn’s multiple comparisons test; *p*=0.2344). In contrast, the mean diameter of medium vessels was 18 ± 4.6 μm in baseline, 23 ± 5.9 μm in induction, and 23 ± 4.9 μm in recovery conditions. The mean diameter of medium vessels in induction and recovery conditions was significantly different compared to baseline conditions, but it was not significantly different between induction and recovery conditions (**Figure 2b**; n = 7 vessels from 4 mice; Kruskal-Wallis test (3, 115.8) and Dunn’s multiple comparisons; *p*<0.0001). Lastly, large vessels had a mean diameter of 38 ± 8.2 μm in baseline, 46 ± 7.4 µm in induction, and 46 ± 4.5 μm in recovery conditions. The mean diameter of large vessels in induction and recovery conditions was significantly different compared to baseline conditions, but it was not significantly different between induction and recovery conditions (**Figure 2c**; n = 4 vessels from 3 mice; Kruskal-Wallis test (3, 82.9) and Dunn’s multiple comparisons; *p*<0.0001).

**Figure 2.**
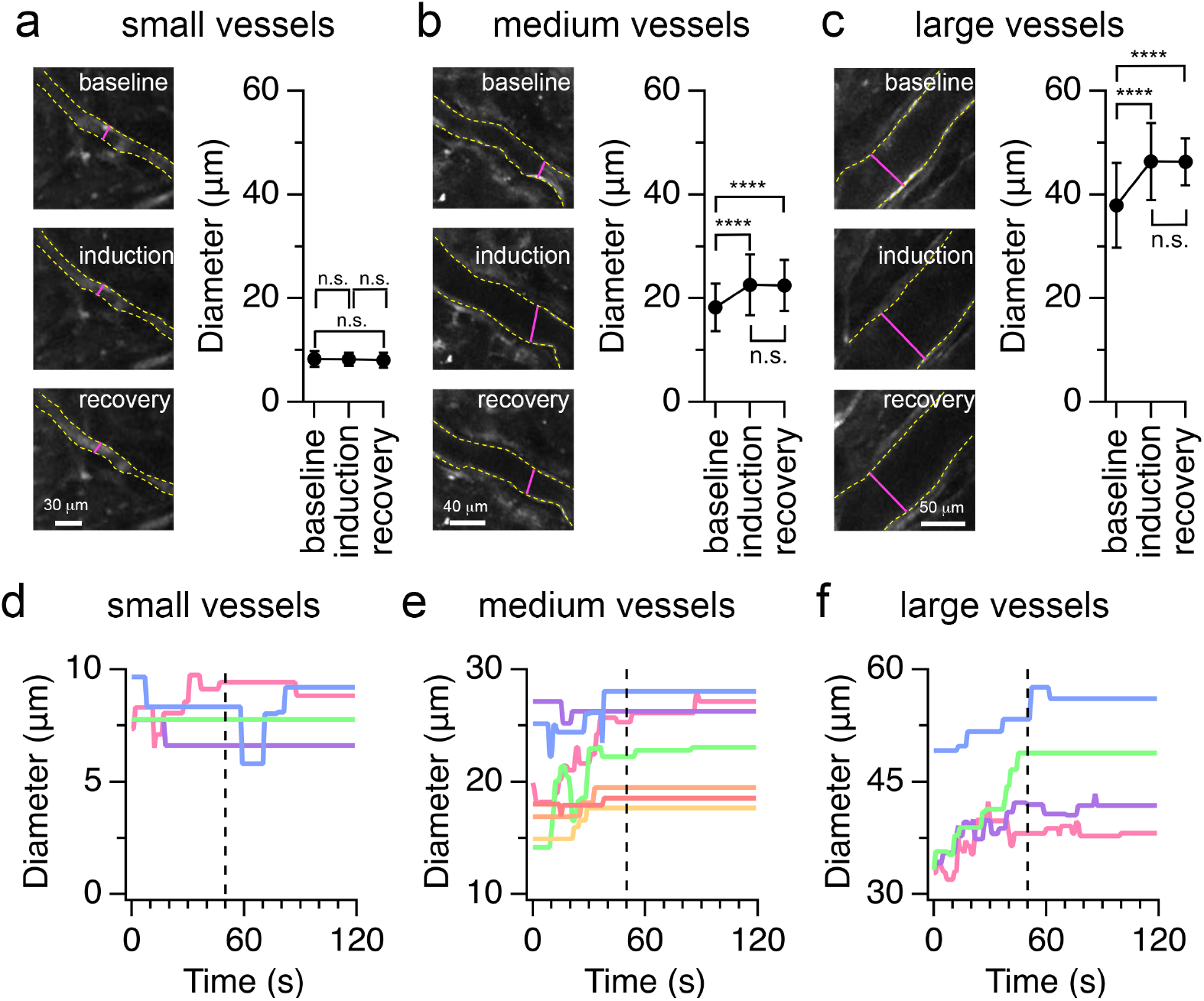
Vasodilation effects of isoflurane (ISO) on vessels of different diameter. **a, b, c**, Exemplar images and diameter plots of small, medium and large vessels in baseline, induction and recovery conditions. Image frames are maximum intensity projections of 60 consecutive images in steady state. Dashed yellow lines show the outline of representative small, medium and large vessels. Magenta lines show the location of length measurement. Diameter data represent mean ± sd (n = 30 replicates per vessel from 4 small vessels; n = 30 diameter replicates per vessel from 7 medium vessels; n = 30 diameter replicates per vessel from 4 large vessels). **d, e, f**, Diameter plots of individual small, medium and large vessels during the induction condition (n = 120 consecutive diameter measurements per vessel). Color indicates different vessels. Vertical dashed line marks the 50 s interval after ISO flow at time zero. Statistical significance was determined with the Kruskal-Wallis test and Dunn’s multiple comparisons, alpha = 0.05; **** = *p*< 0.0001; n.s. = not significant.

To estimate the time to vessel dilation during the induction of anesthesia, we plotted individual vessel diameter as a function of time. This approach showed that the diameter of small vessels did not change systematically after the onset of ISO flow at time zero. In contrast, dilation of the majority of medium diameter vessels and large diameter vessels occurred within a 50 s window, as indicated with vertical dashed lines in **Figure 2d-f**. Altogether, these results demonstrate ISO-induced dilation in medium and large diameter vessels, but not in small diameter vessels.

### ISO inhibits calcium reporter activity in vessels of all sizes

Next, we determined the effect of ISO on endothelial cell calcium reporter activity. As shown in **Figure 3a-c**, we obtained DF/F_0_ recordings from box regions of interest (box ROI) placed on individual vessel segments. With an arbitrary threshold of 3 times the standard deviation (SD) above the mean F_0_, we identified fluorescence increase events of variable duration and amplitude in small, medium and large diameter vessels. Visual inspection of those recordings showed a gradual decrease in DF/F_0_ during the induction condition, and a gradual increase in DF/F_0_ during the recovery condition. To summarize the observed changes in DF/F_0_, we plotted the mean DF/F_0_ in bins of 50 s duration in baseline, induction and recovery conditions (**Figure 3d-f**). In small diameter vessels, mean DF/F_0_ values decreased and increased slightly during the baseline condition. Mean DF/F_0_ values showed a ramp up followed by a ramp down during the induction condition, and a ramp up during the recovery condition (**Figure 3d**). In medium diameter vessels, mean DF/F_0_ values were relatively constant during the baseline condition. Mean DF/F_0_ values showed a clear ramp down during the induction condition, and a ramp up during the recovery condition (**Figure 3e**). Lastly, large diameter vessels showed a ramp up followed by a ramp down during the base line condition, a partial ramp down during the induction condition, and a complex response profile with a ramp up followed by a ramp down during the recovery condition (**Figure 3f**).

**Figure 3.**
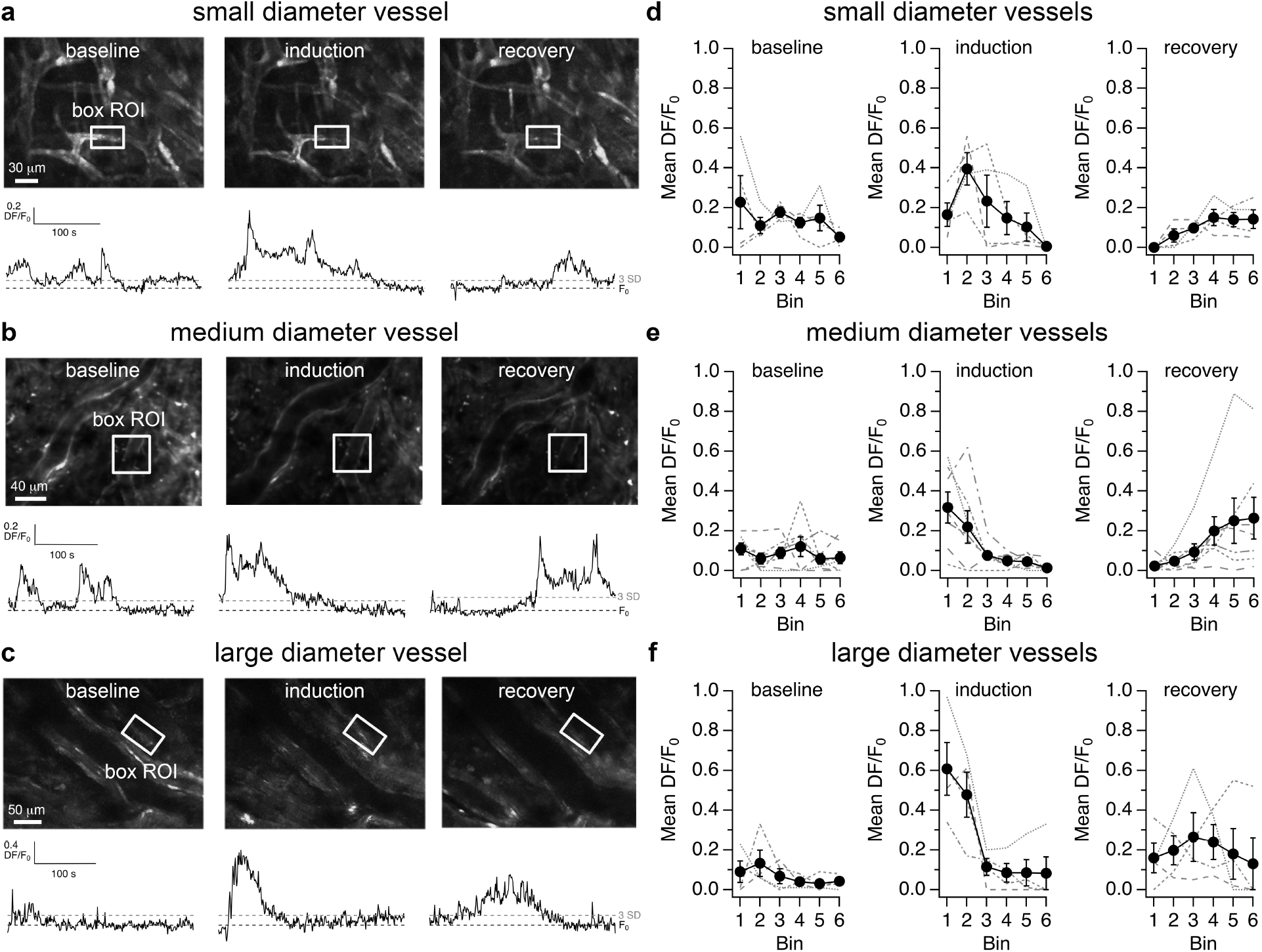
Time course of calcium reporter activity in baseline, induction and recovery conditions. **a, b, c**, Exemplar images and DF/F_0_ recordings measured in regions of interest (ROI) from cerebral blood vessels of different diameter. Black and gray dashed lines indicate mean F_0_ and 3 SD threshold level, respectively. **d, e, f**, Mean DF/F_0_ binned plots of small, medium and large vessels in baseline, induction and recovery conditions. Gray lines represent mean DF/F_0_ for individual blood vessels. Black lines and symbols represent group mean ± sem (n = 4 small vessels; n = 7 medium vessels; n = 4 large vessels). Bin width = 50 s.

To quantify the effects of ISO on calcium reporter fluorescence events, we used normalized DF/F_0_ (_N_DF/F_0_) data and fitting to equation 2 to estimate the time at which the _N_DF/F_0_ reached 50% of its maximum (Bin half = B50; **Figure 4**). We found shorter B50 values for large and medium vessels compared to small vessels during induction conditions (B50 large = 2.3 ± 48×10^−4^ bins = 114 ± 0.24 s, B50 medium = 2.2 ± 0.17 bins = 110 ± 8.6s; B50 small = 4.5 ± 2.0 bins = 224 ± 99s; **Figure 4b**). Furthermore, we found shorter B50 values in large vessels compared to small and medium vessels during recovery conditions (large vessel B50 = 0.45 ± 0.18 bins = 22± 9.1s; medium vessel B50 = 3.5 ± 72×10^−3^ bins = 174± 3.6s; small vessel B50 = 2.0 ±0.79 bins = 99± 39s; **Figure 4d**). In sum, these results show that ISO inhibits calcium reporter fluorescence events in vessels from all sizes. Inhibitory effects were faster in large and medium diameter vessels compared to small diameter vessels. Removal of ISO was followed by a recovery of calcium reporter activity. Recovery time was faster in large diameter vessels compared to medium and small diameter vessels.

**Figure 4.**
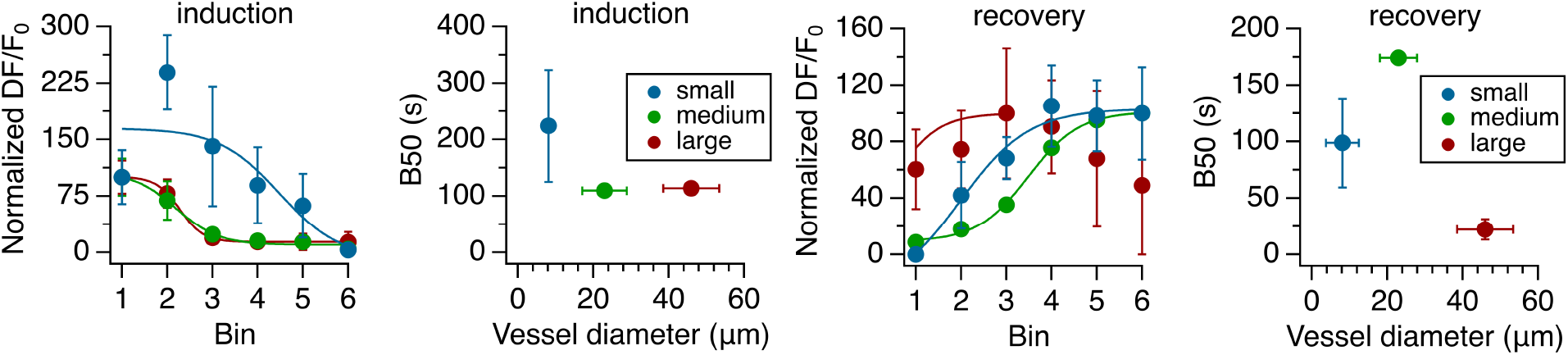
Quantification of isoflurane effects during induction and recovery conditions. **a**, Normalized DF/F_0_binned plots during anesthesia induction. **b**, Plot of B50 values obtained from equation fitting to data in a. **c**, Normalized DF/F_0_binned plots during recovery from anesthesia. **d**, Plot of B50 values obtained from equation fitting to data in c. Data in a and c represent group mean ± sem (n = 4 small vessels; n = 7 medium vessels; n = 4 large vessels). Continuous lines in a and c are fits to equation 2 described in methods. Color code in b and d applies to a and c. Bin width = 50 s.

### ISO disrupts synchronicity of calcium events in vessels of all sizes

Next, we examined DF/F_0_ recordings from multiple small box ROI placed along individual blood vessels of different diameter and compared their time course with respect to DF/F_0_ recordings obtained from the entire image frame (**Figure 5**). In frames where small diameter vessels were imaged, we observed fluorescence increase events that occurred in synchrony and matched maximum peak events observed in frame recordings (vertical black line in baseline, vehicle, induction and recovery conditions in **Figure 5a**). Such synchronous events were observed at a low rate of ∼3.3 mHz. Although frame DF/F_0_ maxima occurred at variable times during baseline and vehicle conditions, we found that during anesthesia induction there was a clear increase of fluorescence events that occurred within 50 s after the onset of ISO flow and that locked around the maximum peak of the frame recording. Of interest to us, synchronous events did not occur during the maintenance condition. Instead, fluorescence events were scattered throughout the recording and did not generate visible peaks in DF/F_0_ frame recordings. Lastly, we observed a gradual increase in synchronous fluorescence events during the recovery condition (**Figure 5a**).

**Figure 5.**
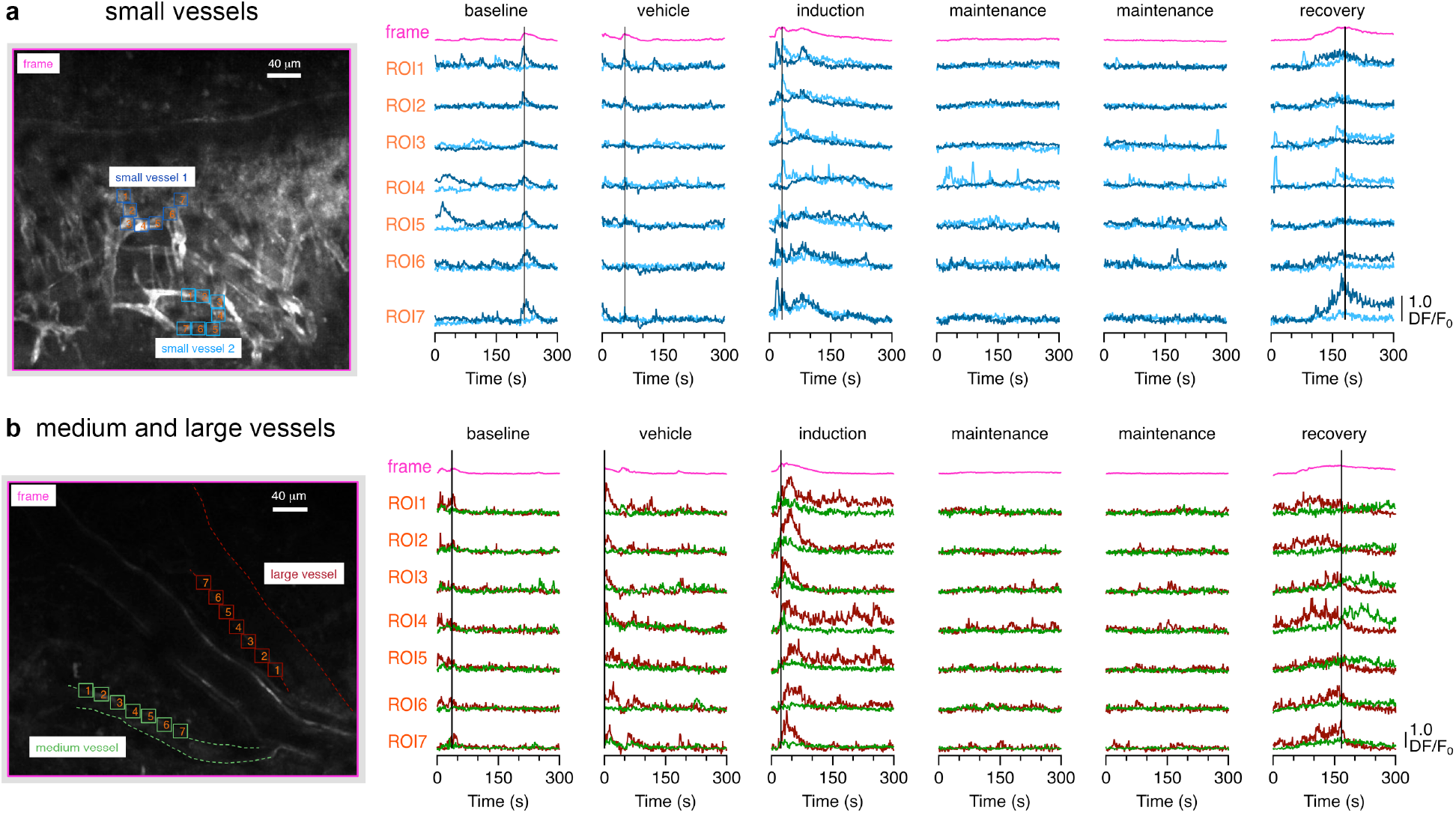
Evidence of synchronous activity in small, medium and large diameter vessels. **a**, Exemplar image frame containing a large diameter vessel (top) and several small diameter vessels (bottom). Seven regions of interest (ROI) were drawn along two small diameter vessels (indicated in dark and light blue). DF/F_0_ recordings from vessel ROI are plotted below the frame DF/F_0_ recording in different conditions. A vertical dashed black line indicates the maximum DF/F_0_ peak in the frame recording for each condition. **b**, Exemplar image frame containing one medium and two large diameter vessels. Seven ROI were drawn along one large diameter vessel and one medium diameter vessel (indicated in dark red and green, respectively). DF/F_0_ recordings from vessel ROI are plotted below the frame DF/F_0_ recording in different conditions. A vertical dashed black line indicates the maximum DF/F_0_ peak in the frame recording for each condition.

Synchrony between frame and box ROI fluorescence events was also observed during baseline and vehicle conditions in recordings from medium and large diameter vessels. Similar locked responses to ISO delivery were observed between DF/F_0_ box ROI and frame recordings during the induction condition (**Figure 5b**). In addition, non-synchronous scattered fluorescence events were observed in the maintenance condition, and a return to synchronous events was observed during the recovery condition. Altogether, these results show that ISO disrupts the occurrence of synchronous fluorescence events in cerebral blood vessels of all sizes.

### The dynamic relationship between changes in vessel diameter and calcium reporter fluorescence is disrupted by ISO

Based on the observations that vessel dilation and fluorescence increase events occurred within a 50 s window after the start of the induction condition, we performed a more detailed analysis of correlated changes in vessel size and calcium reporter activity. In our last analysis, we used polygon ROI that followed the contour of blood vessels to measure simultaneous changes in the area and calcium reporter fluorescence of individual vessels. **Figure 6a-c** shows DA/A_mean_ and DF/F_mean_ traces and scatter plots for small, medium and large diameter vessels, respectively.

**Figure 6.**
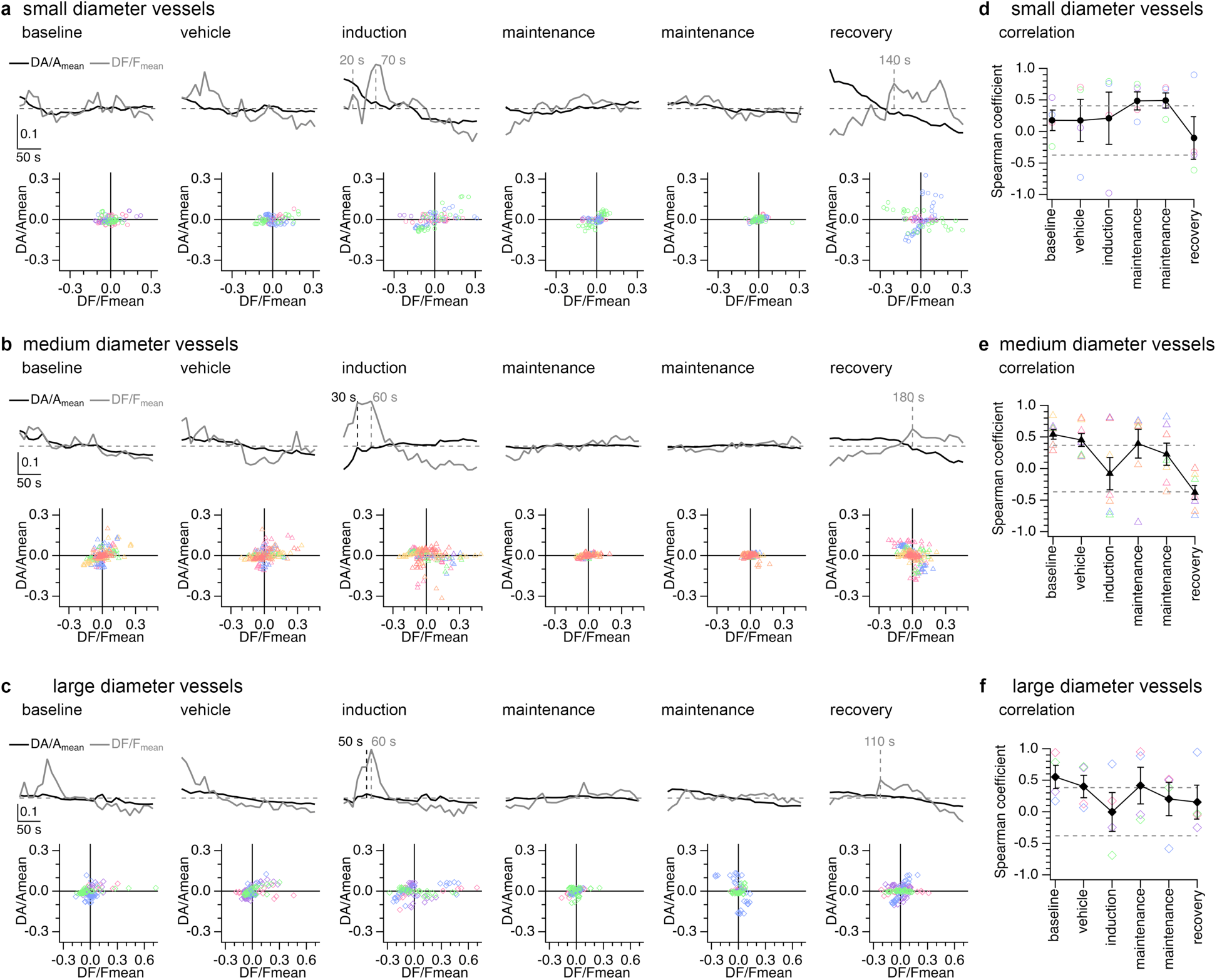
Relationship between vessel size and calcium activity. **a, b, c**, Average DF/F_mean_ and DA/A_mean_ recordings (top) and individual vessel scatter plots (bottom) for small, medium and large vessels in different conditions. **d, e, f**, Spearman correlation coefficients obtained from DF/F_mean_ and DA/A_mean_ scatter plots in small, medium and large vessels in different conditions. Color symbols indicate individual vessels and black lines and symbols represent mean sem (n = 4 small vessels; n = 7 medium vessels; n = 4 large vessels). Dashed horizontal lines represent significance level for positive and negative correlations.

We found that in small vessels, DF/F_mean_ changes had a larger range compared to DA/A_mean_ changes during baseline and vehicle conditions. Furthermore, during the induction condition we confirmed the occurrence of two maxima in the DF/F_mean_ recording at 20 s and 70 s (**Figure 6a**). In agreement with our previous results, we did not observe evidence of vessel dilation during anesthesia induction. However, to our surprise we found that the DA/A_mean_ recording showed a continuous decrease from above A_mean_ values to below A_mean_ values over time, suggesting vessel constriction in response to ISO (see also exemplar images in **Figure 2a**). During the first period of anesthesia maintenance, we found evidence of a slow return from below mean values to mean values in the DF/F_mean_ trace, and a relatively constant DA/A_mean_ trace. During the second period of anesthesia maintenance, DF/F_mean_and DA/A_mean_ recordings showed relatively stable fluctuations. Lastly, we observed a behavior that recapitulated the induction of anesthesia during the recovery condition. While there was a gradual increase in DF/F_mean_ such that the first peak above the mean fluorescence occurred at 140 s, the DA/A_mean_ signal showed a monotonic decrease throughout the recording, suggesting a disruption between vessel tone and calcium reporter activity observed in baseline conditions (**Figure 6a**). To quantify the observed changes, we obtained correlation coefficients from DA/A_mean_ and DF/F_mean_ scatter plots. As a group, small diameter vessels did not show statistically significant correlations in baseline, vehicle and induction conditions. However, there were statistically significant positive correlations during anesthesia maintenance, and a non-significant trend to a negative correlation in the recovery condition (**Figure 6d**).

As expected from our previous results, medium diameter vessels showed different DF/F_mean_ and DA/A_mean_ profiles compared to small vessels. Over the course of baseline and vehicle conditions, there were systematic fluctuations such that fluorescence changes above the mean correlated with area changes above the mean, and fluorescence changes below the mean correlated with area changes below the mean. As a result, statistically significant positive correlations were obtained in grouped data in baseline and vehicle conditions (**Figure 6e**). During anesthesia induction, there were two DF/F_mean_ maxima at 30 s and 60 s, such that they coincided with the first two maxima in the DA/A_mean_ trace. The behavior in the DA/A_mean_ trace was consistent with ISO-induced vessel dilation (**Figure 6b**), and as a result there was a drop in the correlation coefficient to a positive non-significant value (**Figure 6e**). During anesthesia maintenance we observed a non-significant trend to increased positive correlations. Lastly, upon ISO removal DA/A_mean_ showed evidence of constriction, which preceded the first maxima above the mean in the DF/F_mean_ trace (**Figure 6b**), and as a result, there was a statistically significant negative correlation in the group data (**Figure 6e**).

Finally, large diameter vessels had a similar behavior compared to medium diameter vessels (**Figure 6c**), and had statistically significant positive correlations during baseline and vehicle conditions (**Figure 6f**). During the induction condition we found a maximum in the DA/A_mean_ at 50 s and a maximum in the DF/F_mean_ trace at 60 s (**Figure 6c**). We found that correlation dropped to a non-significant value during the induction condition (**Figure 6f**). We note that there was a positive statistically significant correlation during the first period of anesthesia maintenance, and non-significant correlations during the second period of maintenance and the period of recovery (**Figure 6f**).

## Discussion

The relationship between vessel diameter and calcium signaling in endothelial cells in vivo was first studied in peripheral vessels in the context of agonist-induced vasodilation (Tallini et al., 2007). In this study, we used isoflurane (ISO) to induce vasodilation and found two main effects on calcium responses of cerebrovascular endothelial cells (CVECs). There was a short latency increase in calcium activity, followed by a longer latency decrease in calcium activity observed in all vessel types. Since small diameter vessels did not dilate in response to ISO, one may conclude that these changes in calcium activity were independent of vessel dilation. However, the data indicates that the synchrony of calcium events in CVECs was affected by ISO. On one hand, the short latency ISO-induced increase in calcium activity was synchronous within and across different vessels, which is consistent with a global response to ISO exposure and vessel dilation. In agreement with this interpretation, the longer latency ISO-induced decrease in calcium activity was observed in all vessel types, which then exhibited small amplitude desynchronized events during the ISO maintenance condition. Synchronicity of CVEC calcium activity returned slowly during the recovery condition. Furthermore, this slow recovery was not observed for two point measures of vessel diameter, but was noted when measuring changes in vessel area (DA/A_mean_), which suggests that a return to a vessel constricted state correlated with an increase in synchronous calcium activity.

Although we are not aware of previous reports on excitatory effects of ISO on cells of the neuro-glia-vascular unit, there have been numerous studies that demonstrated inhibitory effects of ISO and other general anesthetics on different neural cell types including neurons, astrocytes, mural cells, and microglia (Glück et al., 2021; Schummers et al., 2008; Thrane et al., 2012; Umpierre et al., 2020). Indeed, volatile anesthetics such as ISO, have multiple protein targets, are potent cerebral vasodilators and there exists a narrow range of isoflurane concentrations (1.0 to 1.5%) under which neurovascular coupling is preserved (Masamoto and Kano, 2012; O’Herron et al., 2016; Schlegel et al., 2015; Schummers et al., 2008; Shumkova et al., 2021). Interesting to us, previous reports in auditory brainstem slices showed that ISO reduced action potentials and neurotransmitter release at a half-maximum inhibitory concentration (IC50) of 1.5% (Wu et al., 2004; Wang et al., 2020). The long-latency (∼100 s) ISO-dependent inhibition of CVEC calcium activity observed in this study is similar to that reported recently for ISO-induced inhibition of electrical activity in auditory brainstem neurons measured in neonate rats in vivo (Di Guilmi and Rodríguez-Contreras, 2021). These considerations lead us to speculate that the short latency synchronous increase in CVEC calcium activity may be a direct response to vessel dilation, while the longer latency decrease in CVEC calcium activity and desynchronization could be an indirect response to vessel dilation and ISO diffusion into the brain parenchyma.

In summary, we determined dynamics of ISO exposure on cerebrovascular endothelium calcium activity in awake mice using two-photon imaging through thinned-skull cranial windows. We discovered dual effects of ISO on calcium activity in cerebral vessels with different size. The correlation between vascular tone and calcium activity was disrupted during and after ISO exposure. Although the mechanisms of synchronous and asynchronous calcium activity in endothelial cells were not identified in this study, we propose that synchrony of CVEC calcium activity may be an important feature of the cerebral vasculature that deserves further study. Based on these results we propose that there is a feedback mechanism between intracellular calcium fluctuations in CVECs and the maintenance of cerebrovascular tone.

## Acknowledgements

This work was supported by start up funds (L.S.) and a Harvey L. Karp Discovery Award in the Sciences (A.R.-C.). The mouse strain Cdh5^BAC^-GCaMP8 was developed by CHROMus™, which is supported by the National Heart Lung Blood Institute of the National Institute of Health under award number R24HL120847.

## References

Andresen, J., Shafi, N.I., Bryan, R.M. Jr. (2006) Endothelial influences on cerebrovascular tone. J Appl Physiol; 100(1), 318–27.

Bagher, P., Beleznai, T., Kansui, Y., Mitchell, R., Garland, C.J., Dora, K.A. (2012) Low intravascular pressure activates endothelial cell TRPV4 channels, local Ca2+ events, and IKCa channels, reducing arteriolar tone. Proc Natl Acad Sci USA; 109, 18174–18179.

Bukhari, Q., Schroeter, A., Rudin, M. (2018) Increasing isoflurane dose reduces homotopic correlation and functional segregation of brain networks in mice as revealed by resting-state fMRI. Sci Rep; 8(1), 10591.

Ciobanu, L., Reynaud, O., Uhrig, L., Jarraya, B., Le Bihan, D. (2012) Effects of anesthetic agents on brain blood oxygenation level revealed with ultra-highfield MRI. PLoS ONE; 7, e32645.

Constantinides, C., Murphy, K. (2016) Molecular and integrative physiological effects of isoflurane anesthesia: the paradigm of cardiovascular studies in rodents using magnetic resonance imaging. Front Cardiovasc Med; 3, 23.

Dalal, P.J., Sullivan, D.P., Weber, E.W., Sacks, D.B., Guzer, M., Grumbach, I.M., Brown, J.H., Muller, W.A. (2021) Spatiotemporal restriction of endothelial cell calcium signaling is required during leukocyte transmigration. J Exp Med; 218(1), e20192378.

Di Guilmi, M.N., Rodríguez-Contreras, A. (2021) Characterization of developmental changes in spontaneous electrical activity of medial superior olivary neurons before hearing onset with a combination of injectable and volatile anesthesia. Front Neurosci; 15, 654479.

Dora, K.A., Hill, M.A. (2013) Measurement of changes in endothelial and smooth muscle Ca(2+) in pressurized arteries. Methods Mol Biol; 937, 229–238.

Filippini, A., D’Amore, A., D’Alessio, A. (2019) Calcium mobilization in endothelial cell functions. Int J Mol Sci; 20, 4525.

Gao, Y.R., Ma, Y., Zhang, Q., Winder, A.T., Liang, Z., Antinori, L., Drew, P.J., Zhang, N. (2017) Time to wake up: Studying neurovascular coupling and brain-wide circuit function in the un-anesthetized animal. Neuroimage; 153, 382–398.

Glück, C., Ferrari, K.D., Binini, N., Keller, A., Saab, A.S., Stobart, J.L., Weber, B. (2021) Distinct signatures of calcium activity in brain mural cells. Elife; 10, e70591.

Guerra, G., Lucariello, A., Perna, A., Botta, L., De Luca, A., Moccia, F. (2018) The role of endothelial Ca2+ signaling in neurovascular coupling: a view from the lumen. Int J Mol Sci; 19, 938.

Harris, E.S., Nelson, W.J. (2010) VE-cadherin: at the front, center, and sides of endothelial cell organization and function. Curr Opin Cell Biol; 22, 651–658.

Kansui, Y., Garland, C.J., Dora, K.A. (2008) Enhanced spontaneous Ca2+ events in endothelial cells reflect signaling through myoendothelial gap junctions in pressurized mesenteric arteries. Cell Calcium; 44, 135–146.

Li, C.X., Patel, S., Wang, D.J., Zhang, X. (2014) Effect of high dose isoflurane on cerebral blood flow in macaque monkeys. Magn Reson Imaging; 32(7), 956–60.

Masamoto, K., Kanno, I. (2012) Anesthesia and the quantitative evaluation of neurovascular coupling. J. Cereb. Blood Flow Metab; 32, 1233–1247.

Mumtaz, S., Burdyga, G., Borisova, L., Wray, S., Burdyga, T. (2011) The mechanism of agonist induced Ca2+ signaling in intact endothelial cells studied confocally in in situ arteries. Cell Calcium; 49, 66–77.

O’Herron, P., Chhatbar, P.Y., Levy, M., Shen, Z., Schramm, A.E., Lu, Z., Kara, P. (2016) Neural correlates of single-vessel haemodynamic responses in vivo. Nature; 534, 378–382.

Rothman, J.S., Silver, R.A. (2018) NeuroMatic: An integrated open-source software toolkit for acquisition, analysis and simulation of electrophysiological data. Front Neuroinform; 12, 14.

Schlegel, F., Schroeter, A., Rudin, M. (2015) The hemodynamic response to somatosensory stimulation in mice depends on the anesthetic used: implications on analysis of mouse fMRI data. NeuroImage; 116, 40–49.

Schummers J, Yu H, Sur M (2008) Tuned responses of astrocytes and their influence on hemodynamic signals in the visual cortex. Science; 320, 1638–1643.

Shumkova, V., Sitdikova, V., Rechapov, I. et al. (2021) Effects of urethane and isoflurane on the sensory evoked response and local blood flow in the early postnatal rat somatosensory cortex. Sci Rep; 11, 9567.

Slupe, A.M., Kirsch, J.R. (2018) Effects of anesthesia on cerebral blood flow, metabolism, and neuroprotection. J Cereb Blood Flow Metab; 38(12), 2192–2208.

Stenroos, P., Pirttimäki, T., Paasonen, J., Paasonen, E., Salo, R.A., Koivisto, H., Natunen, T., Mäkinen, P., Kuulasmaa, T., Hiltunen, M., Tanila, H., Gröhn, O. (2021) Isoflurane affects brain functional connectivity in rats 1 month after exposure. Neuroimage; 234, 117987.

Sullender, C.T., Richards, L.M., He, F., Luan, L., Dunn, A.K. (2022) Dynamics of isoflurane-induced vasodilation and blood flow of cerebral vasculature revealed by multi-exposure speckle imaging. J Neurosci Methods; 366, 109434.

Tallini, Y.N., Brekke, J.F., Shui, B., Doran, R., Hwang, S.M., Nakai, J., Salama, G., Segal, S.S., Kotlikoff, M.I. (2007) Propagated endothelial Ca2+ waves and arteriolar dilation in vivo: measurements in CX40BAC-GCaMP2 transgenic mice. Circ Res; 101, 1300–1309.

Taylor, M.S., Francis, M. (2014) Decoding dynamic Ca(2+) signaling in the vascular endothelium. Front Physiol; 5, 447.

Thakore, P., Earley, S. (2019) Transient receptor potential channels and endothelial cell calcium signaling. Comp Physiol; 9(3), 1249–1277.

Thrane, A.S., Thrane, V.R., Zeppenfeld, D., Lou, N., Xu, Q., Nagelhus, E.E., Nedergaard, M. (2012) General anesthesia selectively disrupts astrocyte calcium signaling in the awake mouse cortex. Proc Natl Acad Sci USA; 109(46), 18974–18979.

Umpierre, A.D., Bystrom, L.L., Ying, Y., Liu, Y.U., Worrell, G., Wu, L.-J. (2020) Microglial calcium signaling is attuned to neuronal activity in awake mice. eLife; 9, e56502.

Vestweber, D. (2008) VE-cadherin: the major endothelial adhesion molecule controlling cellular junctions and blood vessel formation. ArteriosclerThrombVasc Biol; 28, 223–232.

Wang, H.Y., Eguchi, K., Yamashita, T., Takahashi, T. (2020) Frequency-dependent block of excitatory neurotransmission by isoflurane via dual presynaptic mechanisms. J Neurosci 40(21), 4103–4115.

Wu, X.S., Sun, J.Y., Evers, A.S., Crowder, M., Wu, L.G. (2004) Isoflurane inhibits transmitter release and the presynaptic action potential. Anesthesiology; 100(3), 663–70.

